# A diffusion model of viral evolution predicts mutation fitness and evolutionary trajectories

**DOI:** 10.64898/2026.09.16.752245

**Authors:** Jinghan Wu, Xiao Ding, Aiping Wu

**Affiliations:** State Key Laboratory of Common Mechanism Research for Major Diseases, Suzhou Institute of Systems Medicine, Chinese Academy of Medical Sciences & Peking Union Medical College, Suzhou, Jiangsu, China, 215123

## Abstract

Viral evolution arises from random mutations and natural selection, yet computational approaches rarely model these two forces in a unified way. We present Viral Evolution Simulator (VES), a diffusion model-based framework that mirrors this duality by design: forward noise injection simulates stochastic mutation, and reverse denoising recapitulates selective filtering. Trained solely on viral protein sequences, VES predicts mutational fitness without functional data, measuring fitness as the reconstruction difficulty of a mutated sequence relative to its wild-type counterpart. Across immune escape, receptor binding, and deep mutational scanning datasets, VES outperforms state-of-the-art generative models, achieving a 31.78% error reduction over the best baseline in immune escape mutation fitting evaluation. When trained on sequences collected before June 2024 and evaluated against H1N1 strains that later emerged, VES assigned high scores to 16 of 20 mutations that subsequently showed the sharpest frequency shifts. Extending to avian influenza H5, the framework reveals a dynamic interplay between antigenic escape and human-type receptor binding. Both functions dropped sharply in 2021, followed by a sustained rise in receptor affinity that could connect to recent epidemiological trends. VES offers a generalizable, sequence-only foundation for tracing evolutionary trajectories and prioritizing mutations for surveillance and experimental validation, pointing toward where functional efforts might matter most.

---

Viruses undergo rapid and continuous genetic variation when they spread. Under the pressure of natural selection, mutations that facilitate infection and transmission are preserved, thus driving the viral evolution [1, 2]. Some of these mutations can significantly enhance the ability to infect human hosts and evade host immunity, leading to further transmission and global spread, posing a major threat to global health. Therefore, studying and predicting viral evolution provide an essential basis for controlling viral spread. However, the inherent randomness and multi-factorial complexity of this process make it highly challenging to accurately model and predict viral evolution. With the advancement of sequencing technologies and the accumulation of genomic data, research on viral evolution has primarily relied on phylogenetic methods to establish viral lineages and reconstruct evolutionary histories [3–6]. Nevertheless, these approaches can only reconstruct evolutionary history based on existing sequence data. They offer limited insight into the molecular mechanisms underlying viral evolution and cannot anticipate its future trajectory.

To bridge this gap between retrospective evolution reconstruction and prospective mutation prediction, increasing attention has been paid to machine learning and deep learning algorithms. For example, Hie *et al*. developed an evolutionary velocity framework based on a protein language model and likelihood gradients to predict the directionality of protein evolution [7]. Similarly, MLAEP model combined a protein language model with multi-task learning to simulate the directed evolution trajectories of the viruses using sequences and structural data of the receptor binding domain [8]. More recently, E2VD integrated functional priors with deep neural networks to identify and predict the key mutation drivers during the evolutionary process [9]. Although these models achieve strong performance in modelling viral evolution, they generally rely on supervised learning with functional data to impose function constraints, and overlook the coupled effects of stochasticity and determinism in the viral evolution [10, 11]. As a result, their capacity to capture fine-grained evolutionary dynamics and to generalize to unseen selective pressures remains limited.

Generative models offer a fundamentally different paradigm. Rather than learning a discriminative mapping from sequence and structural data to viral functions, they model the probability distribution over sequence space, thereby capturing both the deterministic evolution constraints and stochastic sequence variation within a unified framework. In a seminal study, DeepSequence employed a variational autoencoder (VAE) trained on protein sequences in an unsupervised manner, demonstrating that the latent structure of natural sequence distributions inherently encodes the functional constraints and accurately predicting the fitness effects of mutations across diverse protein families [12]. Building on this idea, the EVEscape frame-work incorporated immune escape-related constraints into the VAE-based fitness calculation to predict the escape potential of viral mutations [13].

Among generative approaches, diffusion models have emerged as one of the most powerful frameworks, achieving remarkable success across diverse research fields in the life sciences [14, 15]. Notably, diffusion models can themselves be viewed as evolutionary models, sharing a compelling mechanistic analogy with biological evolution [16]. The forward noising process mirrors the stochastic accumulation of mutations, whereas the reverse denoising process parallels the action of natural selection, providing a predictive basis for potential evolutionary trajectories. Leveraging this analogy, DiffEvol employed reverse diffusion dynamics to decipher viral evolution constraints, reconstructing the manifold space of viral genotypes to demonstrate viral evolution process [17]. While DiffEvol provided interpretable computational constraints on viral evolution, it did not extract molecular-level evolutionary features or offer mechanistic insights, and thus could not evaluate or predict the functional consequences of potential evolutionary trajectories, substantially limiting its generalizability and practical utility. More broadly, we posit that viral evolution is a directed process in which numerous random mutations are shaped by complex natural selection. The evolutionary process therefore cannot be captured by any single biological constraint, but must instead be characterized through the interplay of multiple constraints.

Therefore, in this work, we propose Viral Evolution Simulator (VES), an unconstrained diffusion model-based framework for simulating and predicting viral evolution. Trained solely on viral protein sequences, VES learns the natural distribution of viral sequence space and predicts mutations with high fitness in nature populations, substantially reducing the reliance on large-scale functional assays. Across extensive experimental datasets spanning multiple viruses, VES shows strong agreement with measured mutational fitness. When partial functional measurements are available, VES can further incorporate simple functional constraints to predict the functional consequences of viral evolution, providing computational support for the proactive prevention and control of emerging infectious diseases.

## Results

### VES: Characterizing Viral Evolution Dynamics via Diffusion Framework

Viral evolution can be conceptualized as a process of mutation diffusion [18], in which the continuous generation of random mutations introduces noise into the genetic space of the genome sequences, while functional selection restricts the direction of viral evolution, shaping its evolutionary trajectory [19, 20]. Consistent with these principles, diffusion models mirror the stochasticity of random mutation and the determinism of natural selection through noise diffusion and denoising procedures [16]. Specifically, the forward process incrementally injects Gaussian noise into the data, analogous to the continuous accumulation of viral mutations, whereas the reverse denoising process reconstructs data from a chaotic distribution. This reverse process conceptually recapitulates the filtering mechanism of natural selection, demonstrating immense potential for simulating and predicting evolutionary trajectories (Fig. 1a). Grounded in this mechanistic convergence, we introduce Viral Evolution Simulator (VES), a diffusion model-based paradigm that quantitatively describes the viral evolutionary process and natural fitness landscapes. In contrast to existing methods that use diffusion processes to describe viral evolution [17], VES employs an unconstrained diffusion model to learn latent evolutionary features from historical viral sequences, enabling the simulation and prediction of advantageous mutations at the molecular level and providing a versatile computational foundation for applications such as vaccine strain recommendation and cross-species transmission prediction. Furthermore, compared with generative models such as VAE and pre-trained transformer in viral language models [21], the close alignment between the diffusion mechanism and the viral evolution process reduces the demand for diversity beneath viral sequence data, as well as the large amount of training data.

**Fig. 1.**
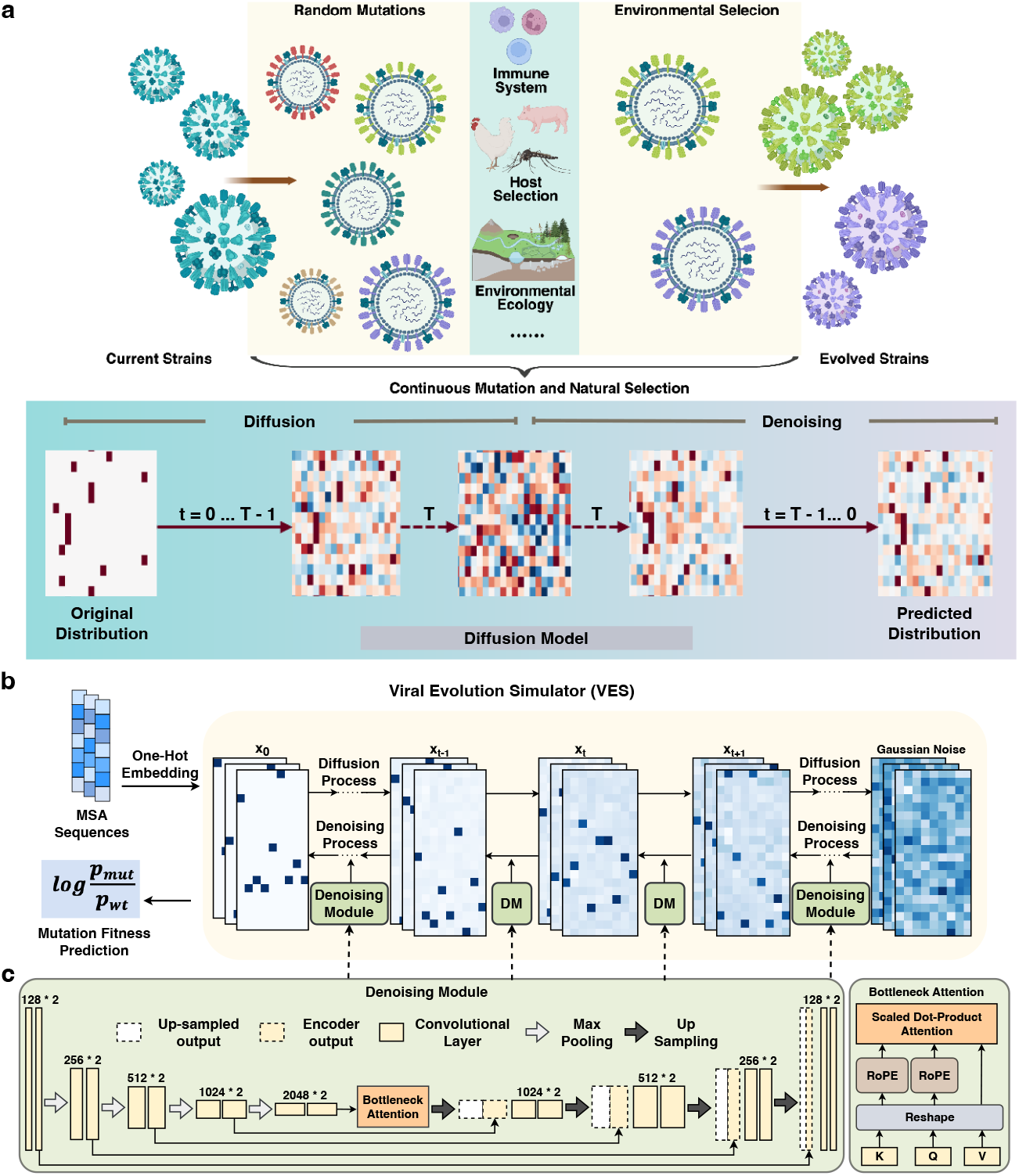
Schematic overview of the Viral Evolution Simulator (VES) **a**, Conceptual analogy between viral evolution and diffusion-based sequence modelling. Top, viral populations evolve from current strains to new strains through continuous mutation and natural selection. Bottom, the corresponding procedures 0 of diffusion models, in which the original sequence distribution is progressively corrupted by Gaussian noise during the forward process (*t* = 0 to *T*) and reconstructed through iterative denoising during the reverse process (*t* = *T* to 0), yielding the predicted distribution. **b**, Workflow of VES. Sequences from multiple sequence alignment (MSA) are one-hot encoded (x_**0**_), diffused towards Gaussian noise (x_**T**_) in the forward process, and reconstructed by denoising modules (DM) in the reverse process. The log-likelihood ratio between a mutant sequence and its wild-type counterpart, denoted as 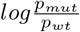, is derived from the reconstruction likelihoods and serves as the predicted mutational fitness. **c**, Architecture of the denoising module. A modified U-Net comprising five convolutional blocks with two convolutional layers each (channel dimensions indicated) is implemented, with an expanded input channel configuration (128 channels), and a bottleneck attention block integrated with rotary position embedding (RoPE, right inset) 9 to capture long-range dependencies between sites across viral protein sequences.

In implementation, VES utilized the one-hot encoded representations of input multiple sequence alignments (MSAs) to capture continuous, high-dimensional evolutionary features from extant strains, and subsequently reconstructs these input embeddings from the noisy distribution via a specialized denoising module. A simplified version of Denoising Diffusion Probabilistic Model (DDPM) [22] diffusion paradigm was adopted to extract latent evolution features. Crucially, during the reverse denoising process, we defined the log-likelihood ratio of a mutated sequence relative to its wild-type counterpart as the quantitative metric for natural evolutionary fitness, as proposed in previous work [12]. This ratio fundamentally reflected the comparative difficulty of reconstructing mutant versus wild-type sequences. A higher reconstruction barrier indicated that the mutation deviated significantly from the learned empirical distribution of evolutionary sequences, thus statistically less favoured to survive under natural selection (Fig. 2b).

**Fig. 2.**
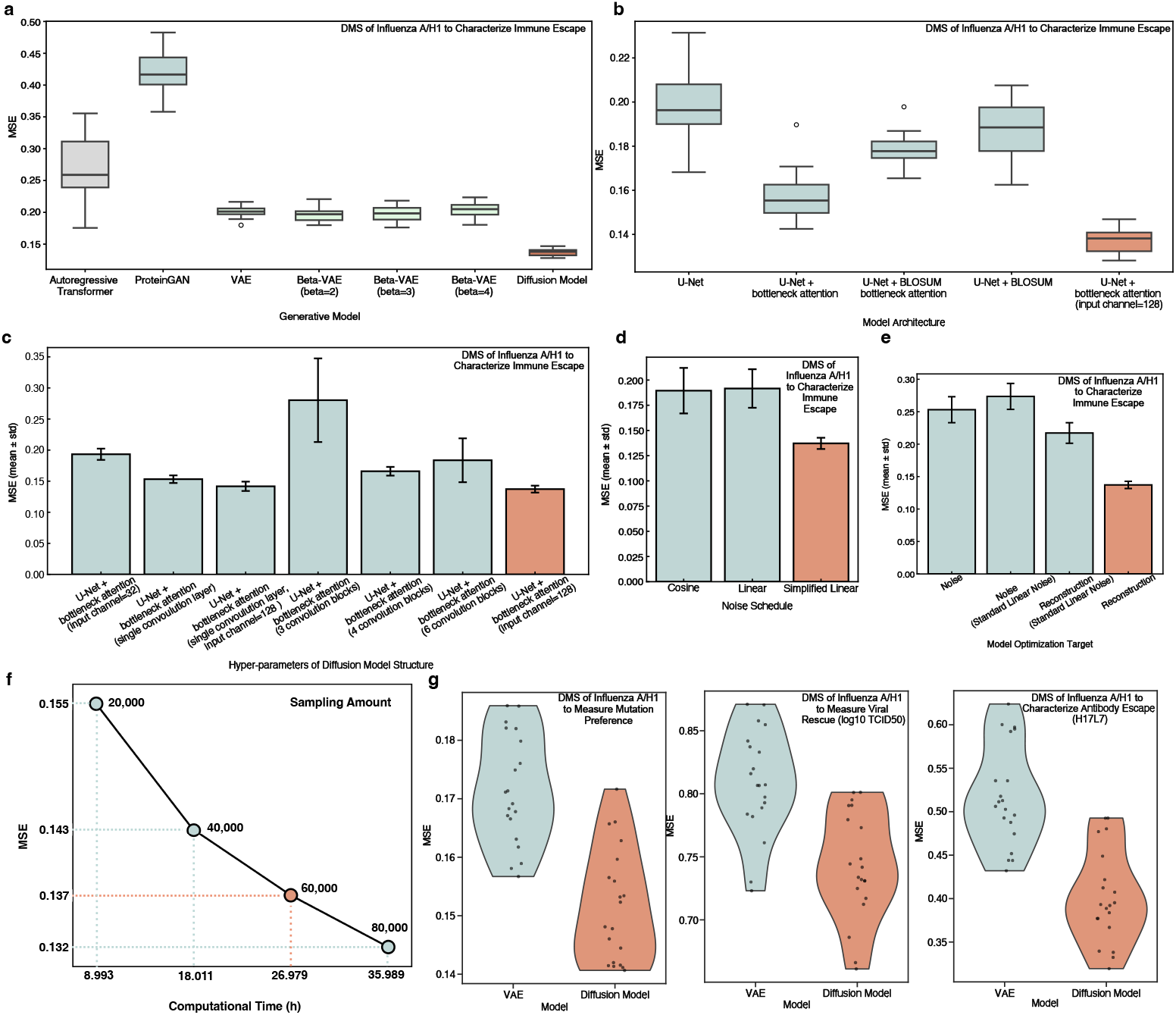
Quantitative benchmarking and ablation analyses of VES. **a**, Benchmarking of mutational fitness predictions against experimentally measured immune escape profiles. Boxes show the distribution of mean squared error (MSE) across 20 Monte Carlo repeated subsampling tests for the autoregressive transformer, ProteinGAN, VAE and three beta-VAE variants, as well as VES (Diffusion Model) on the influenza A/H1 DMS dataset to characterize immune escape. **b-e**, Ablation of VES components on influenza A/H1 DMS immune escape data, quantifying the individual contributions of denoising module architecture (b), input channel width and convolutional depth (c), noise schedule (d) and training objective (e). Bars show *mean ± std* across 20 independent runs. The full VES configuration is shown in orange. **f**, Trade-off between Monte Carlo sampling volume, prediction error (MSE) and computational time (hours) on the influenza A/H1 DMS dataset to characterize immune escape. 60,000 trajectories (orange) are selected for all subsequent analyses. **g**, Generalization to varying DMS tests for functional measurements, including mutation preferences (rescaled), viral rescue experiments and antibody escape (H17-L7), comparing VES with the VAE baseline. Violins show the distribution of MSE across 20 independent runs, with individual data points overlaid.

Tailored to the intrinsic global characteristics of viral protein sequences, we adapted the classic backbone U-Net architecture [23] to comprise five convolution blocks, each containing two convolutional layers, for evolutionary simulation. Specifically, we incorporated a bottleneck attention block integrated with Rotary Position Embedding (RoPE), which substantially enhanced the capacity of VES for long-range dependency modelling, thereby effectively capturing non-local epistatic site dependencies within natural evolutionary trajectories. Furthermore, we expanded the number of input feature channels from the commonly used 64 to 128 to facilitate the extraction of high-dimensional evolutionary representations. This architectural refinement yielded substantial performance gains in our representation and reconstruction evaluations (Fig. 1c and Fig. 2c). Comprehensive implementation details of VES are provided in Methods.

### VES Outperforms Existing Generative Models in Quantitative Comparison

To evaluate the predictive robustness of VES, we benchmarked its performance against several state-of-the-art generative paradigms, including VAE [13] with its variants, autoregressive transformer [24, 25] and ProteinGAN [26], across diverse viral species (Fig. 2, Extended Data Fig. 1). Particularly, we adopted the same VAE model used in EVEscape [13], currently the most widely used and highly competitive generative model in viral evolution research. We first evaluated all models on deep mutational scanning (DMS) datasets of viral immune escape compiled from prior research [13]. Rather than applying an arbitrary binary threshold to classify escape mutations, we directly correlated the raw probability outputs of the model with continuous experiment measurements to circumvent potential threshold-selection biases. Among the evaluated benchmarks, VES achieved the lowest Mean Squared Error (MSE) of 0.1372 and smallest standard deviation (*σ* = 0.0056), demonstrating a 31.78% improvement (*P <* 0.0001, Monte Carlo subsampling test) over the VAE-based framework employed in EVEscape that previously established as a top performer [13]. It yielded an MSE of 0.2011 (*σ* = 0.0094), which nonetheless outperformed the Autoregressive Transformer and ProteinGAN frameworks (Fig. 2a). VES maintained this advantage on SARS-CoV-2 DMS profiles, achieving the lowest MSE of 0.1431, albeit with a slightly higher variance potentially driven by the intrinsic heterogeneity of the underlying clinical data cohorts (Fig. 2a, Extended Data Fig. 1).

To delineate the orthogonal contributions of our architectural innovations, we conducted a systematic ablation study. The empirical evaluation revealed that while a baseline U-Net diffusion model naturally surpassed the VAE benchmark with an MSE of 0.1978, the integration of our bottleneck attention mechanism substantially enriched long-range sequence modelling, further driving the MSE down to 0.1574 for influenza virus datasets (Fig. 2b). Scaling the input convolutional layer from the standard 64 feature channels to 128-channel configuration optimized performance, consistent with the capacity of VES to exploit high-dimensional evolution landscapes (Fig. 2c). Notably, architectural optimization was context-dependent, and a streamlined convolutional topology with fewer blocks suited SARS-CoV-2 representations, contrasting with the deeper hierarchy required for influenza (Extended Data Fig. 1c). Beyond structural modifications, refining the diffusion generative process also played an essential role in improving model performance, specifically implementing a simplified noise schedule and a reconstruction-focused training objective that was consistently mitigated prediction errors (Fig. 2d and Fig. 2e). Given that VES employed Monte Carlo sampling to approximate the log-likelihood ratios for mutation fitness, we also calibrated the sampling volume. A trajectory count of 60,000 struck the optimal trade-off, delivering minimized MSE with acceptable computational latency, whereas 40,000 samples suffered from under-sampling errors and 80,000 diminished computational efficiency without significant accuracy gains (Fig. 2f).

Beyond immune escape profiles, we generalized our framework across orthogonal experimental dimensions featuring deep mutational preferences, host receptor-binding affinities, and antibody evasion potencies. Across all validated functional landscapes, VES consistently yielded a markedly reduced MSE compared to the VAE baseline. Most notably, VES achieved strong predictive agreement with mutation preference datasets, with an MSE of 0.1516 (*σ* = 0.0097), reflecting the actual evolution constraints in nature, firmly validating its efficacy as a robust and comprehensive simulator for natural viral evolution dynamics (Fig. 2g).

### VES Maps Fitness Landscapes and Viral Evolution Trajectories

As a generalized viral evolution simulator, VES shows a strong ability to simulate and predict adaptive fitness in viral evolution. We leveraged high-throughput DMS datasets to evaluate its capability in identifying and predicting dominant variants for seasonal influenza A/H1 (Fig.3) and A/H3 (Extended Data Fig. 2) subtypes [27, 28]. Mechanistically, the mutational fitness landscape simulated by VES displayed a distinguished topological congruence with the baseline DMS mutation tolerability across individual residues in HA1 domain. Notably, this included residues 145 and 179 (132 and 166 in H3 numbering), which represent critical, lineage-defining sites underlying the antigenic drift of post-2009 pandemic H1N1 strains (Fig. 3a). The distributions of model predictions and DMS measurements showed substantial overlap, with Jensen-Shannon divergence of 0.0179 and Bhattacharyya coefficient of 0.9774 (Fig. 3b). The predictions of VES also captured the highly adaptive mutations of the influenza A/H3 viruses, though the overall distribution was more concentrated than that of the DMS data (Extended Data Fig. 2a,b).

**Fig. 3.**
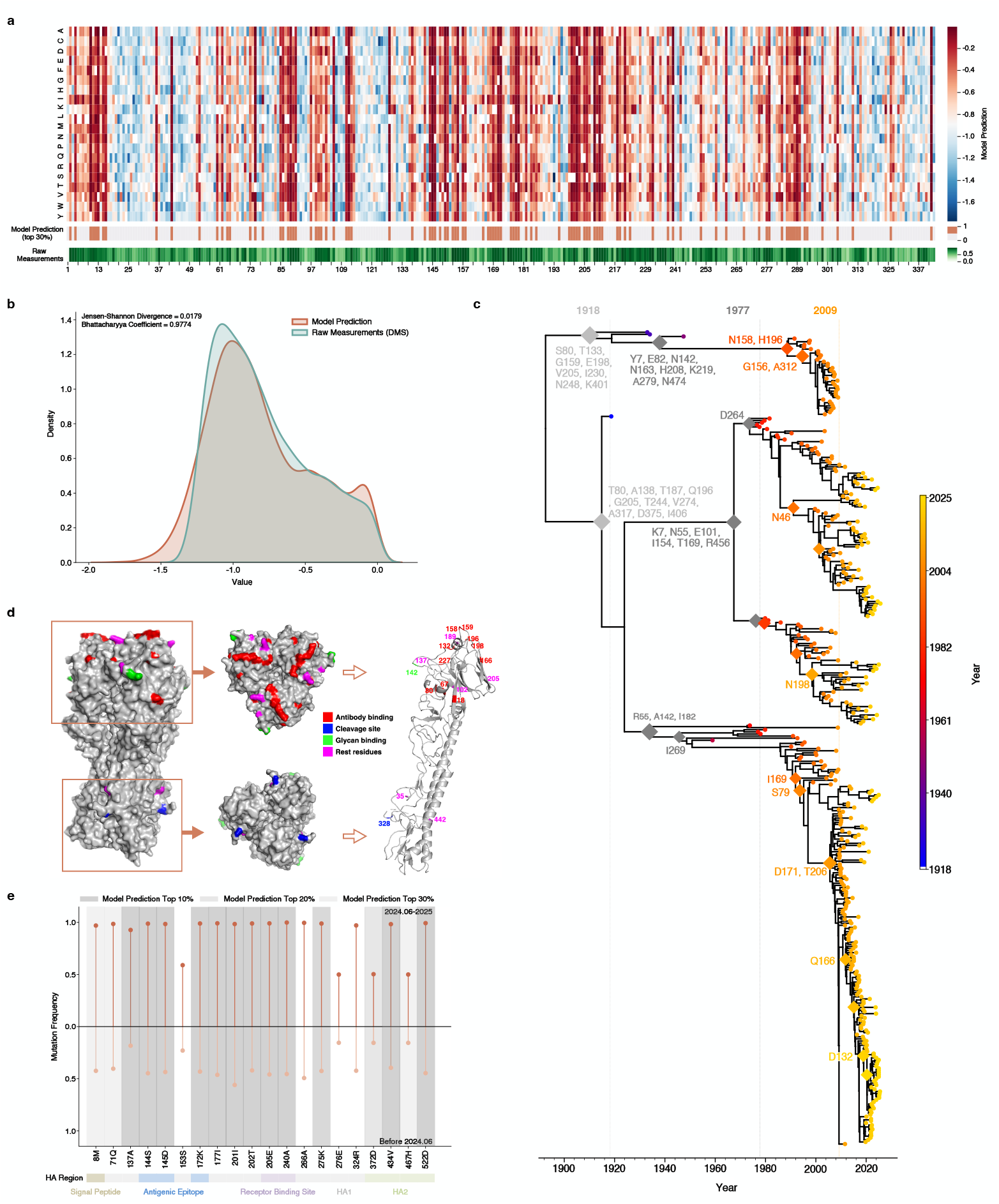
VES maps mutational fitness landscapes and evolutionary trajectories of influenza A/H1 viruses. **a**, Heatmap of site-specific mutational fitness predicted by VES across the HA1 domain. Top-ranked sites prioritized by the model and experimentally validated functional sites are indicated below. **b**, Density distributions of experimentally measured DMS fitness values and VES-predicted mutational fitness, showing substantial agreement (Jensen–Shannon divergence of 0.0179 and Bhattacharyya coefficient of 0.9774). **c**, Time-calibrated phylogeny of influenza A/H1 viruses, with high-scoring mutations (top 5% of VES predictions) mapped onto the tree, spanning the major evolutionary stages of seasonal A/H1. **d**, Structural mapping of the top 20 high-fitness sites predicted by VES onto the haemagglutinin (HA) monomer (PDB: 1RVX). **e**, Prospective validation against H1N1 sequences circulating after June 2024 and entirely excluded from training: 16 of the 20 mutations with the largest post-cutoff frequency shifts ranked within the top 30% of VES predictions, 12 of which ranked in the top 10%.

Beyond capturing advantageous mutations and sites at the single-strain level, VES also simulated the dynamic evolutionary process of seasonal influenza viruses since the beginning of their surveillance. We constructed the time-calibrated phylogenetic tree to comprehensively characterize the evolutionary trajectories of seasonal influenza A/H1 and A/H3 viruses and identified lineage-specific mutations at key internal nodes. Mutations whose predicted fitness scores fell within the top 5% were retained and mapped onto the phylogenetic tree. Among these sites, predictions for residues 200 and 91 (187 and 82 in H3 numbering) ranked within the top 30 across the entire protein, aligning with historical evidence that alterations at these sites significantly influenced the emergence and prevalence of distinct viral lineages. The high-scoring mutations predicted by VES encompassed the entire evolutionary process and key evolutionary stages of seasonal influenza viruses, providing a computational foundation for elucidating the molecular dynamics of viral evolution (Fig. 3c and Extended Data Fig. 2c).

### VES Provides Computational Priors for Viral Evolution Prediction

To assess the practical utility of VES for predicting viral evolutionary trends, we examined whether its per-site fitness predictions could serve as reference priors for experimental studies. Given that DMS assays pinpointed the immunogenic globular head of H1 HA as a hypertolerant zone for mutations [27], VES faithfully replicated this biological reality, and assigned peak fitness landscapes to residues at key structural sites. Specifically, we selected the top 20 sites predicted by VES and mapped their positions on the HA protein monomer. These sites covered functionally important regions, including antibody-binding sites, N-linked glycosylation sites, the signal peptide, and the cleavage sites, 14 of which have been validated in prior experimental studies (Fig. 3d, Extended Data Fig. 2d, and Extended Data Table 1, Table 2). Crucially, VES also captured the distinct evolutionary pattern of the H3 subtype, which exhibited elevated mutational tolerance within the stalk domain and a more homogenous tolerance profile across the HA structure relative to H1 [28]. VES accurately recovered high-scoring sites of influenza A/H3 viruses at both the distal boundaries of the HA head (e.g., site 205 in H3 numbering) and the lower stalk architectures (e.g., sites 377 and 466 in H3 numbering), regions historically implicated in antibody binding and in determining the pH threshold of membrane fusion [29, 30] (Extended Data Fig. 2d).

Beyond characterizing established evolutionary patterns, VES can forecast mutations with future epidemic potential. We evaluated VES against independent H1N1 sequences circulating after June 2024, which were entirely excluded from the training cohorts. Utilizing MAFFT [31], these circulating sequences were aligned against the historical baseline to track directional selection. By filtering for residues exhibiting the most distinguished frequency shift between the pre- and post-June 2024 epochs, we isolated 20 emerging specific mutations. Remarkably, 16 of these observed mutations were recovered within the top 30% of VES fitness predictions, with 12 ranking in the top 10% (Fig. 3e). These results indicate that high-scoring mutations predicted by VES carry information about potential epidemic spread, providing a computational basis for the proactive surveillance of emerging strains.

Importantly, VES provided not only broad evolutionary trajectories, but also a basis for more specialized functional analyses. Natural selection in reality is shaped by specific, often competing functional constraints. For instance, the high mutational tolerance observed in the immunogenic head largely reflects the interplay of host immune escape and receptor-binding constraints. By incorporating targeted functional constraints into the diffusion process, VES can be further tailored to probe finer-grained functional dimensions of viral evolution. Leveraging this capability, we next extended VES to predict the evolution of specific phenotypes, including antigenic drift and receptor-binding affinity.

### VES Models Antigenic Drift and Immune Escape

For rapidly evolving viruses, antigenic drift and immune escape are key determinants of long-term circulation and population-level fitness. Therefore, accurately anticipating these changes is critical for understanding viral adaptation and informing surveillance of potential epidemics. As shown in the Fig. 4a, positions within antigenic epitopes that received high model-derived fitness scores also exhibited elevated probabilities of antibody escape. When epitope-specific constraints were incorporated into the diffusion process, the distribution of model predictions closely approached the distributions measured by DMS assays across multiple neutralizing antibodies (Fig. 4b).

**Fig. 4.**
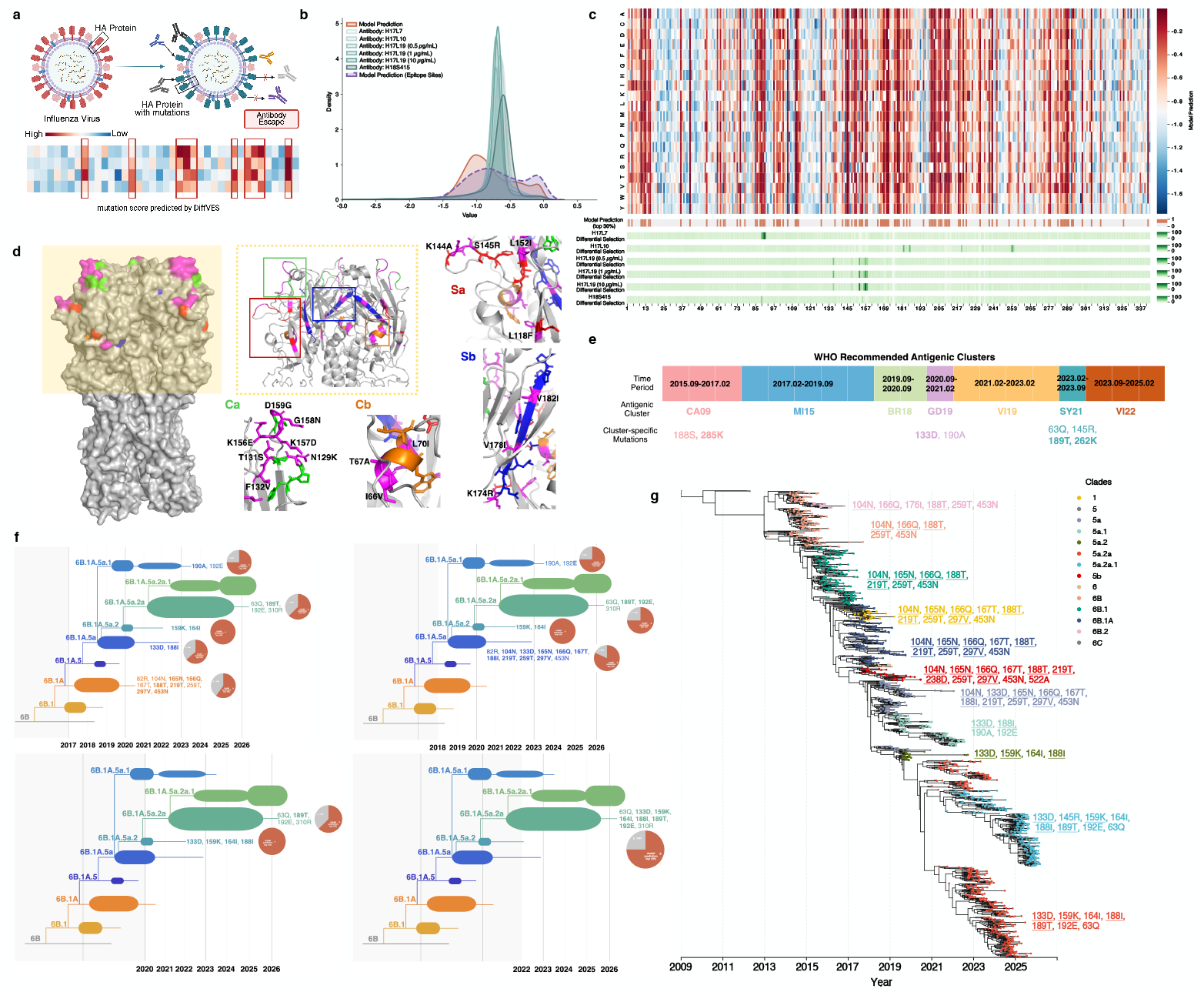
VES predicts antigenic evolution and clade dynamics under functional constraints. **a**, Schematic of incorporating epitope-specific constraints into the VES framework for antibody escape prediction. **b**, Density distributions of experimentally measured antibody escape profiles (DMS) compared with VES fitness predictions generated under epitope constraints, across multiple neutralizing antibodies. **c**, Heatmap of site-specific mutational scores across the HA1 domain under epitope constraints, with residues ranked in the top 30% of model predictions and experimentally validated escape sites indicated below. **d**, Structural mapping of VES predictions onto the haemagglutinin (HA) monomer, with canonical epitope regions of influenza A/H1 shown in distinct colours and mutational sites in the top 10% of VES predictions highlighted in magenta. **e**, Prediction of antigenic cluster-defining mutations. Half of the mutations defining the antigenic clusters represented by WHO-recommended vaccine strains over the past decade ranked within the top 5% of VES predictions, and all remaining mutations, except 201S (188S in H3 numbering), for which the reference sequence already encoded the cluster-defining residue, ranked within the top 30%. **f**, Retrospective prediction of clade evolution in influenza A/H1. VES was trained on sequences collected strictly before each of four time points (2017, 2018, 2020 and 2022), ensuring that subsequent clade dynamics were entirely unseen during training. Pie charts show the pro-portion of clade-specific mutations, which were prevalent after each cut-off, ranked within the top 10% of predictions. Results for influenza A/H3 (2014) are shown in Extended Data Fig. 4. **g**, Phylogenetic tree of influenza A/H1 lineages (Nextstrain) with dominant mutations (top 30% of VES predictions) mapped onto clade branches. Mutations ranked in the top 5% are underlined, while clade-specific mutations of the prevailing 5a.2a.1 lineage and its D.3 subclade are highlighted in light blue.

Residues ranked within the top 30% of the model predictions accurately mapped onto immunodominant sites previously validated in neutralizing antibody selection experiments [32]. These included key residues 153, 157, and 158 (140, 144, and 145 in H3 numbering) targeted by antibody H17-L19, residues 182 and 186 (169 and 173 in H3 numbering) for antibody H17-L10, residues 89, 90, and 91 (80, 81, and 82 in H3 numbering) for H17-L7, and residue 155 (142 in H3 numbering) for antibody H18-S415 (Fig. 4c). Mapping these predictions onto the three-dimensional structure revealed that surface-exposed patches of the hemagglutinin (HA) protein, which were critical for paratope-epitope interactions, consistently overlapped with high-scoring predictions (top 10%), indicating a pronounced susceptibility to localized immune evasion. This pattern was consistently recapitulated when analyzing avian influenza A/H5 HA profiles. VES identified heightened antibody escape potentials within both the globular head and the upper stalk architectures, capturing several experimentally characterized variants, including R90K and S125bN (H3 numbering) [33] (Fig. 4d and Extended Data Fig. 3c).

Existing experiments have focused on a limited number of antibodies and genetic backgrounds, leaving most of the mutational space, particularly the mutations defining emerging antigenic clusters. In contrast, VES is not restricted by the choice of experimental strains or antibodies. It employed evolutionary computation to reveal the process of antigen evolution and can characterize antigenic evolution at the molecular level, including transitions between antigenic clusters. We examined the antigen clusters represented by WHO-recommended vaccine strains over the past decade and selected representative mutations defining each cluster. Half of these cluster-defining mutations received scores within the top 5% of VES predictions with the exception of 201S (188S in H3 numbering), which was the wild-type amino acid used in model calculation. The evolution predictions for the rest cluster-specific mutations all fell in the top 30% range (Fig. 4e) as well. Thus, VES not only reproduces known escape events, but also predicts which mutations may drive viral evolution towards emerging antigenic clusters.

### VES Identifies and Predicts Potential Epidemic Trends in Clade Dynamics

Building on the ability of VES to forecast antigenic clusters, we further extended this capability to the recognition and prediction of clade-specific mutational trajectories. We defined the mutations ranking in the top 30% of the model predictions as dominant mutations in viral evolution, and mapped them to the phylogenetic tree provided by Nextstrain [5]. These mutations were evenly distributed across clade branches at distinct evolutionary stages, and most of them were potential adaptive advantage mutations with high model prediction values (the underlined mutations in the Fig. 4g, top 5% in model prediction values). Since June 2024, circulating human seasonal A(H1N1) pdm09 viruses have been dominated by HA clade 5a.2a.1, with increasing predominance in the 2024-2025 season. Within this clade, D.3 has emerged as a representative subclade of 5a.2a.1 [34]. VES successfully identified its clade-specific mutations (marked in light blue), among which four mutations possessed top 5% model predictions. Subclade-specific mutations 120A and 372V [34, 35] also obtained top 5% and top 30% model prediction scores, respectively, highlighting the potential of VES in exploring the evolution of genetic space.

Furthermore, VES demonstrated the capacity to predict clade-specific mutations before they become prevalent, providing reliable forecasts regarding potential outbreaks. We conducted a retrospective analysis in which we selected four key time points in influenza A/H1 viruses circulation over the past decade, i.e., 2017, 2018, 2020, and 2022, as well as one key time point for influenza A/H3 viruses (2014, Extended Data Fig. 4). For each time point, VES was trained using sequence data collected strictly prior to that year to derive mutation scores. Figure 4f illustrates the clade information and clade-specific mutations prevalent following each retrospective validation year, with pie charts displaying the proportion of clade-specific mutations falling within the top 10% of model predictions. In all retrospective tests, VES assigned high prediction scores to potentially advantageous mutations prior to the corresponding clades becoming prevalent, highlighting its robust capability to predict emerging epidemiological trends, and its generalizability to periods during which selective pressures shift substantially.

### VES Anticipates Human-Type Receptor Binding Adaptation in Viral Evolution

Beyond predicting antigenic evolution, we incorporated receptor-binding site (RBS) constraints into the model predictions, and applied VES to assess the potential capability of avian influenza viruses to bind to human-type receptors (Fig. 5a). Although the distribution of experimental *α*2-6 binding measurements was relatively narrow, largely owing to the limited availability of data across most sites, the distribution of VES prediction scores spanned its upper range and effectively captured mutations with high experimental values, indicating strong potential for identifying substitutions associated with increased *α*2-6 receptor binding affinity. Furthermore, when sequences from a particular host were selected from the training data, VES could be adapted to host-specific model, with the distribution of model predictions shifting towards the experimental results (Fig. 5b,c).

**Fig. 5.**
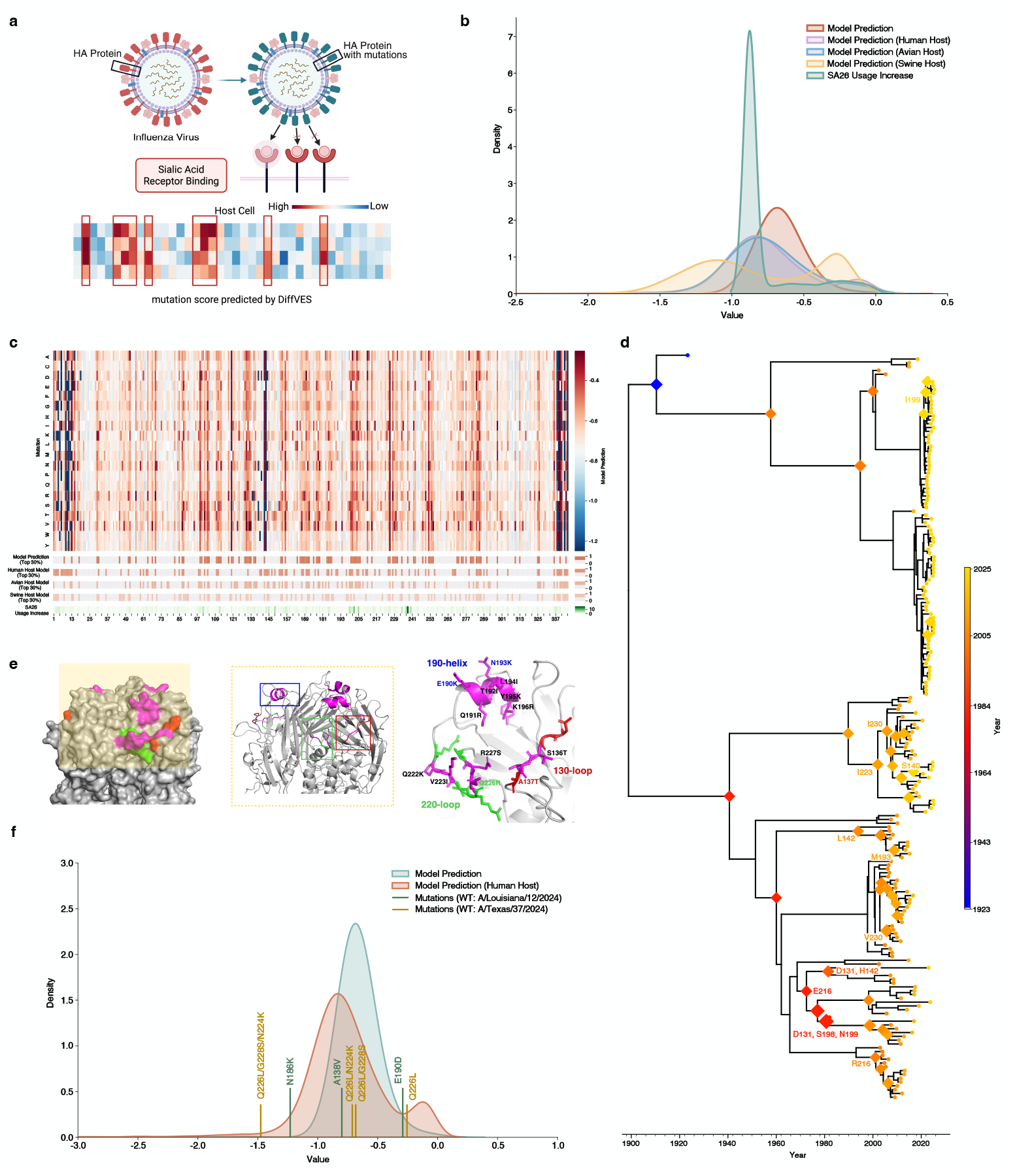
VES forecasts receptor-binding adaptation in H5 avian influenza viruses. **a**, Schematic of incorporating receptor-binding site (RBS) constraints into VES for receptor-binding prediction. **b**, Density distributions of experimentally measured *α*2-6 receptor-binding values and VES prediction scores, together with predictions from host-specific VES models trained on sequences from the selected host, showing a shift towards the experimental distribution. **c**, Site-specific prediction landscape across the HA1 domain of influenza A/H5 HA. High-scoring sites within the RBS and experimentally characterized receptor-binding sites are indicated below. **d**, Time-calibrated phylogeny of influenza A/H5 viruses with high-scoring mutations in the RBS (top 30% of model predictions) mapped onto the tree, spanning the evolutionary history of A/H5 since 1997. **e**, Structural mapping of high-scoring receptor-binding mutations onto the HA protein, with the 130-loop, 190-helix and 220-loop shown in different colours and mutations ranked in the top 10% of predictions highlighted in magenta. **f**, Distribution of VES predictions for mutations with experimentally confirmed enhancement of human-type receptor binding, compared with the overall prediction distribution.

In addition to identifying mutations that enhance human receptor-binding affinity, VES reproduced key mutations that emerged during the evolution of avian influenza viruses over time. Mutations in the receptor binding region associated with high model predictions spanned different evolutionary periods of the influenza A/H5 subtype, especially after 1997, when an H5N1 avian influenza virus was first confirmed to infect humans in Hong Kong [36], reflecting a potential evolutionary trajectory by which avian influenza viruses crossing the species barrier to infect humans (Fig. 5d).

The ongoing epizootic of influenza H5N1 viruses in poultry and wild birds sporadically spills over to mammals, raising an urgent requirement to recognize the mutations that can enhance their binding affinity to human-type receptors during evolution. VES provides a computational foundation for addressing this need and demonstrates great potential for generating experimental priors before receptor binding experiments are conducted. Mutations with top 10% prediction scores were observed across most receptor binding sites, and several of them had been implicated in increased binding to human-adapted receptors, including Q238R, A149T, E202K and N205K (corresponding to Q226R, A137T, E190K and N193K in H3 numbering) [33]. VES accurately prioritized these mutations, assigning three of them to the top 5% of prediction scores. A mutation that significantly increases the ability of avian influenza viruses to bind human-adapted receptors, i.e., Q238L (Q226L in H3 numbering) [37], also ranked within the top 30% of model predictions (Fig. 5e).

### VES Reveals the Molecular Basis Underlying the Co-evolution of Viral Antigenicity and Receptor Binding Affinity

The functional evolution of influenza viruses does not proceed independently. Rather, it but involves complex mechanisms of co-evolution, for instance, with among functions such as antigenicity and receptor affinity. Mutations that enhance binding affinity for human receptors may exhibit varying degrees of evolutionary fitness due to their impact on antigenicity (Fig. 5f). Building on its function-prediction capability, VES can be further utilized to elucidate the co-evolution dynamics between antigenicity (antibody escape) and human receptor binding during the transmission of H5 subtype avian influenza viruses (Extended Data Fig. 5). More than 70% of the mutations with high potential fitness (ranking in the top 30% model predictions) indicated an increase in antibody escape ability, about 30% of which also showed an increase capability to bind with human receptors (Extended Data Fig. 5a). Specifically, antigenic epitopes of HA located near the receptor binding site (RBS) were prone to simultaneously change both antigenicity and receptor-binding capability (Extended Data Fig. 5b). Next, we filtered mutations that obtained high model prediction and frequently emerged within the year as the dominant mutations, and collected the corresponding data from functional experiments, to track the evolutionary trends of both antigenicity and receptor affinity simultaneously. In general, these two functions exhibited an alternating trend of rising and falling over time. However, both functions experienced a sharp decline in 2021, followed by a simultaneous increase the next year. In particular, the receptor binding capability increased significantly, and this high receptor binding affinity persisted until 2025. This may help to explain the molecular basis of influenza virus epidemics in the post-COVID-19 era (Extended Data Fig. 5c). VES alleviates the constraints imposed by functional experimental data on computational models in existing work, facilitating the functional analysis of the evolution of rare or emerging viruses. Moreover, by incorporating functional data, VES can be further used to uncover latent co-evolutionary features across different functions, thereby providing deeper insight into the complex mechanisms of viral evolutionary dynamics.

## Discussion

In this work, we present VES, an unsupervised diffusion model-based framework that learns the natural distribution of viral sequences and quantifies mutational fitness without functional supervision. Across deep mutational scanning benchmarks spanning multiple viruses, VES outperformed state-of-the-art generative models, including VAE-based frameworks, autoregressive transformers and ProteinGAN. The per-site fitness predictions of VES recapitulated experimentally validated functional sites across antigenic, receptor-binding and other structural regions, and captured lineage-defining mutations throughout the evolutionary history of seasonal influenza. In a strictly prospective evaluation, VES recovered 16 of the 20 mutations with the largest frequency shifts in H1N1 populations after June 2024 within the top 30% of its predictions, underscoring its potential for the proactive surveillance of emerging variants. By further incorporating targeted functional constraints, VES resolved the co-evolution of antigenicity and receptor-binding affinity, revealing population-level signatures of shifting selective pressures.

A fundamental question underlying the application of deep learning to viral evolution research is not merely whether a model performs well, but why it is principled to do so. We propose that VES provides a mechanistically grounded answer to this question. The algorithmic correspondence with viral evolution allows VES to implicitly encode both the stochastic and deterministic components of evolutionary dynamics within a unified generative frame-work, without requiring pre-defined fitness information or considerable sequence diversity as prerequisite constraints. This addresses a key limitation of prior computational approaches, which often depend on large, diverse multiple sequence alignments or experimentally annotated fitness conditions that are difficult to collect for many viruses with limited genomic surveillance, thereby constraining their modelling capacity for viral sequences in practice. Unconstrained by functional limitations, VES not only provides a computational foundation for elucidating the molecular mechanisms of viral evolution, but also offers priors on mutational sites for functional experiments from the whole-genome genetic space, demonstrating broad application potential.

Nevertheless, we also recognize certain limitations in present work. First, while we employ diffusion models to extract continuous, high-dimensional evolutionary features from viral genomic sequences, VES learns a time-independent fitness prior. The model itself does not perceive the temporal stage of viral evolution, and temporal resolution is achieved only by training on sequences collected before a defined cut-off. Consistent with this property, when predicting epidemic clades for influenza A/H3 viruses in years not covered by the training data, the coverage of clade-specific mutations by high-scoring mutations gradually decreases as the prediction horizon extends (Extended Data Fig. 4), quantitatively illustrating the expected decay of the time-independent prior under cumulative antigenic drift. Second, existing experimental data for specific functions (such as receptor affinity) remains insufficient, and high-throughput experimental data suffer from distribution bias due to the specific viral strains or antibodies selected for the experiments. Accordingly, the higher variance observed in the SARS-CoV-2 benchmarks, the narrower prediction distribution for A/H3 relative to DMS measurements, and the tendency of *α*2-6 binding predictions to concentrate in the upper range of experimental values may all partly reflect such data limitations rather than intrinsic model behaviour. We further note that the optimal architecture was context-dependent, shallower topologies favoured SARS-CoV-2, whereas deeper hierarchies were required for influenza, which may reflect differences in sequence conservation and alignment depth across viruses and warrants systematic investigation. Some advantageous mutations and residues predicted by VES have not been confirmed by existing experimental results. However, given their location in critical regions, they may still exert a potentially significant influence on viral evolution (Fig. 5c, Extended Data Fig. 2d). Third, the prediction of generative models rely on empirical distributions. Significant shifts in selective pressure can cause substantial divergence of mutations from historical distributions and the evolutionary features learned by the model. Consequently, the prediction of fitness for mutant strains may be constrained, necessitating the incorporation of additional biological priors to enhance predictive performance.

Looking forward, beyond its current scope, the generalizability of VES enables consideration at both the function and species levels. In terms of functional generalizability, VES can be extended to additional phenotypes critical at distinct stages of the viral life cycle, such as replication fidelity and drug resistance governed by polymerases and proteases, virion stability modulated by matrix and nucleoproteins and immune antagonism mediated by non-structural proteins. To achieve this breadth of functional coverage, experimental data including DMS measurements and pseudovirus neutralization assay results can be introduced into VES framework, directly mapping from sequence variation to quantitative functional readouts across diverse viral protein families. In terms of species generalizability, VES has been validated across three influenza subtypes (H1, H3 and H5) as well as SARS-CoV-2, demonstrating an ability to learn evolutionary principles beyond any single virus. We anticipate its extension to a broader spectrum of clinically important viruses such as RSV and HIV, where computational capacity remains limited despite rapidly accumulating surveillance data. More broadly, the conceptual correspondence between noise diffusion-denoising and the stochastic–selective dynamics of molecular evolution suggests that VES could serve as a general framework for modelling adaptive molecular evolution under multidimensional selective pressures, from viral evolution to bacterial adaptation under antibiotic selection.

## Methods

### Data Acquisition

#### Training data

We collected amino acid sequences of viral surface proteins from the Global Initiative on Sharing All Influenza Data (GISAID) database [38] to generate the training dataset in this work. To be specific, we retrieved and downloaded all amino acid sequences of influenza virus hemagglutinin protein available up to May, 2024 on GISAID. After performing quality control and 100% redundancy removal, we retained a total of 95,560 sequences, with 565 amino acids in length and composed exclusively of the 20 standard amino acids for each sequence. Regarding the SARS-CoV-2 viruses, we retrieved and downloaded the spike protein amino acid sequence data available up to June, 2025 on GISAID. After performing quality control and 100% redundancy removal, we retained only those sequences that were 1,273 residues in length and composed exclusively of the 20 standard amino acids, resulting in a total of 55,246 sequences. After dataset generation, we performed multiple sequence alignment using MAFFT, selecting the Influenza A virus (A/WSN/1933(H1N1)) strain and the hCoV-19/Wuhan/WIV04/2019 (WIV04) strain of SARS-CoV-2 viruses as the wild type reference sequences for the respective alignments.

#### Evaluation data

We collected large amount of experimental data on various subtypes of influenza and SARS-CoV-2 viruses, particularly the high-throughput DMS data, to validate the effectiveness of the model prediction. For quantitative evaluation, we primarily adopted the same DMS data [39] used in previous work [13] for a fair comparison on the prediction of immune escape. Besides, we also employed the experimental measurements of mutation preference [27], mutation function changes in receptor binding site [40], and antibody escape [32] in the quantitative fitting evaluation to comprehensively demonstrate the outstanding modelling performance of VES. For the following model prediction analysis of VES, we concentrated on the fitness landscape of influenza A/H1 and A/H3 subtype viruses provided by DMS experiments [27, 28] and the fitting results of mutation fitness prediction given by VES. Then, we further investigated antigenic epitope sites of influenza A/H1 subtype and avian influenza A/H5 subtype viruses [32, 33], as well as the mutations in receptor binding sites of avian influenza A/H5 subtype viruses [33] to evaluate the functional change prediction ability of VES with corresponding DMS data.

To demonstrate the model generalization across viral species, we also employed the same experiential data used in EVEscape model evaluation [13] upon SARS-CoV-2 viruses. Furthermore, we concentrated on the antibody escape and ACE2 binding experimental data [41, 42] to evaluate the function change prediction performance of VES.

### Model Framework of VES

#### Evolutionary simulation and feature extraction using VES

The dynamic process of viral evolution involves the emergence of distinct mutations during different time periods, thus influencing and changing the evolutionary trajectory. From this perspective, we target the simulation of dynamic evolution of viruses, identifying and predicting the dominant mutations across the entire evolution process. Given a dataset of amino acid sequences derived from viral surface proteins, denoted as 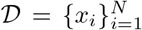, where *x*_*i*_ ∈ ℝ^*N ×Dim×Seq*^ represents the one-hot embedding of amino acid sequences with embedding dimension *Dim* of 20 and sequence length *Seq* of 565, the objective is to construct the high-dimensional feature space *z*_*i*_ and the distribution of naturally existing sequences *p*_*θ*_ with *D*, denoted as: *f*_*θ*_ : *x*_*i*_ → (*z*_*i*_, *p*_*θ*_). We adopt diffusion model to learn the distribution of viral sequences and construct the latent space *f*_*θ*_, making corresponding adjustments to better suit the viral sequence and evolution characteristics.

Compared to the image generation tasks, in which diffusion model shows great success, we implement a simplified version of Denoising Diffusion Probabilistic Model (DDPM) to tackle sequence data. Originally, diffusion model aims to estimate the conditional expectation of *z*_*i*_:

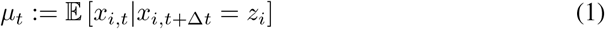

where *t* refers to the time step in the diffusion process, and *µ*_*t*_ is the function at time step *t*. In the training phase, functions across all time steps are estimated by optimizing the denoising objective:

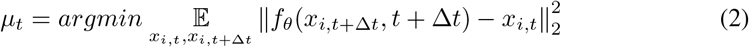

Thus, in the reverse sampling process, we can approach 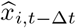 by:

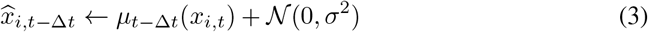

From this perspective, the expectation of *x*_*i,t™*Δ*t*_ is predicted with the trained diffusion model. However, in practice, we predict the expectation of original input *x*_*i*,0_ instead, to reveal the reconstruction difficulty for the mutated sequences, which can be viewed as the distribution difference between mutations and the viral protein sequences in nature. Furthermore, we use the shared model parameters throughout time steps, rather than learning independent functions *µ*_*t*_. Based on the diffusion and denoising framework, we put forward VES to model and simulate the viral evolution process, providing quantitative representations for viral mutations upon model predictions.

### VES architecture

The denoising module of VES is designed upon the classic U-Net model with down-sampling and up-sampling convolutional blocks. For the convolutional layers used here, the kernel size is set to 3 with 1 padding size. To better extract the high-dimensional evolution feature from amino acid sequences, we increase the number of input channels in the first down-sampling block to 128, which is 64 in normal situations. For the down-sampling blocks, two convolutional layers with layer normalization are implemented, with conditional modulation which can be used to introduce restrictions such as time step into the model. Max pooling layer is also attached after convolution, with a kernel size of 2. We utilize five down-sampling blocks, and the number of input feature channels in each block is double that of the preceding one. For the up-sampling blocks with symmetric structure, we use up-sampling layers and convolutional layers to reconstruct the input sequence embedding gradually, and include the conditional modulation for option. Considering the long term dependencies and the interaction among amino acid sites, we further utilize a bottleneck attention block to extract sequence feature. We use rotary position embedding here to maintain the relative positions of amino acid sites in the genome sequences, mapping the site dependencies into geometric structure space when down-sampled into lower dimension. To be specific, the frequency is defined as:

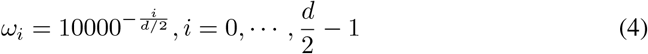

where *d* is the feature dimension. Then, we obtain the sinusoidal embedding:

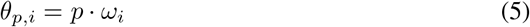

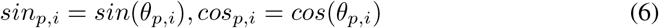

where *θ*_*p,i*_ is the angle of position *p*. For the query 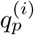 and key matrixes 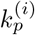 of position *p*, we use rotation transformation for position embedding:

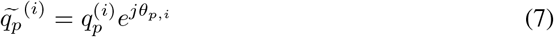

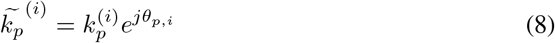

Now, the attention score *attn* is computed as:

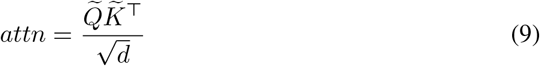

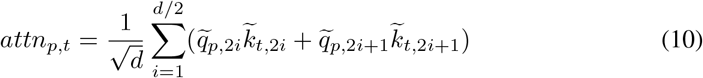

The implementation of rotary position embedding-based bottleneck attention improves the modelling of inter-site interaction, enhancing the global feature extraction ability of VES.

### Mutation fitness prediction

Innovated by the fitness metric computed in EVEscape model, we quantitatively evaluate the mutation fitness in nature through model reconstruction of the input sequence embedding. It is assumed that mutations with higher fitness are more likely to survive under natural selection, and they shall possess similar distribution with wild type, which is smaller difference in the reconstruction loss. Therefore, we define the mutation fitness prediction as the log ratio of the mutation sequence distribution to the wild type given model parameters trained on existing viral sequences, as shown below:

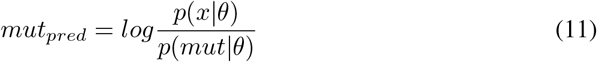

where *θ* refers to the model parameters trained on viral surface protein amino acid sequence data, and the negative log probabilistic can be obtained by:

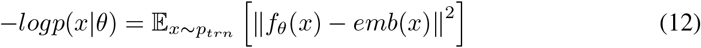

where *p*_*trn*_ stands for the training dataset, and *emb*(*x*) is the embedding of the input sequence. Thus, we turn the evaluation of viral mutation fitness in nature to the calculation of viral evolution modelling, and provide an efficient quantitative representation to predict the evolution outcome of potential mutations.

### Model training objective and strategies

In implementation, the diffusion process is defined as:

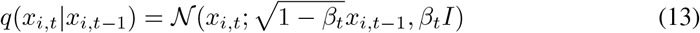

where *β*_*t*_ refers to the intensity of noise at time step *t*. We adopt a simple linear noise schedule to keep the stability in model training with the embedding of amino acid sequence data, which shows a better prediction performance compared to the cosine schedule used in most occasions. Since we predict the expectation of original input sequence embedding given denoising output, the optimization objective of VES is set to minimize the negative log-likelihood of mentioned embeddings:

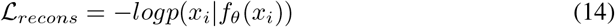

Through the denoising and reconstruction, VES learns the distribution of mutated sequences that can be preserved in entire natural selection, and building the latent evolution feature space.

The model training and evaluation were conducted on the single NVIDIA GeForce RTX 5090 GPU. The AdamW optimizer with learning rate of 1×10^−4^ was employed for model training. Gradient norm clipping and exponential moving average (EMA) were also implemented to prevent gradient explosion during the training of the diffusion model, stabilizing the training procedure.

### Model Evaluation

#### Baseline models

We adopt three generative models and their variants as baseline models for the comparison of prediction performance with VES, i.e., variational autoencoder (VAE), beta-VAE, generative adversarial network (GAN), and autoregressive transformer. They show state-of-the-art generative capability in a wide range of applications, such as computer vision, protein design, and viral function prediction, making them competitive approaches for VES.

#### Variational autoencoder (VAE)

VAE shows successful implementation in the prediction of diseases and viral escape function in previous work, remaining the commonly used model for the natural distribution construction of viral sequence data. In this work, we exactly followed the VAE model architecture and parameters supplied by EVEscape [13] for a fair comparison, and adopted beta VAE, a crucial variant of VAE to further explore its distribution learning ability. We chose three settings of beta, i.e., 2, 3 and 4, to show a thorough comparison of modelling effect between VAE and proposed VES.

#### Generative adversarial network (GAN)

Serving as the classic generation model, GAN also obtains outstanding performance in the bioinformatics fields, such as single-cell RNA-seq data augmentation [43], drug molecular generation [44], protein engineering [45] and so on. Among them, ProteinGAN [26] shows great success in the construction of natural protein sequence space and enables the generation of functional sequences, making it a direct competitor of our model. Therefore, we employed the original implementation of ProteinGAN, and making corresponding adjustments on the calculation of mutation fitness prediction. In particular, we used the discriminator to evaluate difference of distribution between the mutation and wild type, as made quantitative predictions based on it. Firstly, we randomly sampled a batch of natural sequences *o* and put into the discriminator, obtaining the discrimination score *D*(*o*) for the natural distribution. Then, we used the mutated sequences as the input of discriminator to calculated the discrimination score *D*(*mut*), after which the mutation fitness could be evaluated by the difference between above scores:

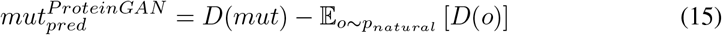

where *p*_*natural*_ referred to the distribution of natural sequences.

#### Autoregressive transformer

Autoregressive transformer is one of the most classic generation models used as language models [46, 47]. In this work, we adopted the basic structure of autoregressive transformer with the 6-layer and 8-head transformer decoder. The input dimension was set to 256, and we revised the size of vocabulary to 21 to include the gap of MSA sequences in tokenization.

#### Ablation study

We made comprehensive comparisons of model architecture, model hyper-parameters, and some of the key factors in the diffusion model to demonstrate the superior performance of VES, revealing how data characteristics affected the model design and implementation performance.

#### Model architecture

We adopted the basic U-Net model with one-dimension convolutional layers and input feature channel of 64 as the backbone of VES in the beginning of this work, illustrating the viral function prediction performance compared to VAE models. Then, we further introduced bottleneck attention module to improve the long-term dependency modelling ability of VES. We also adopted BLOSUM matrix as additional restriction utilizing backbone model, and added it to the viral sequence to verify whether prior knowledge of protein amino acids was beneficial for the extraction of viral evolution patterns.

#### Model hyper-parameters

We further evaluated different model hyper-parameter settings to obtain the optimal viral evolution simulation performance. To be specific, we made changes on the number of convolutional layers in each convolution block (i.e., single layer and double layers), the number of input feature channels (i.e., 32, 64 and 128), as well as the number of convolution blocks (i.e., 3, 4, and 5 blocks) based on our backbone U-Net model.

#### Noise schedule

We evaluated both standard cosine noise schedule, which was raised in improved DDPM [48] and commonly used in diffusion models, and the linear noise schedule in the original DDPM [22]. The cosine noise schedule could be defined as:

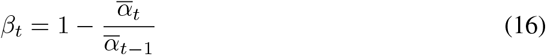

where 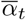 referred to the noise intensity accumulated over time steps, and it could be obtained by:

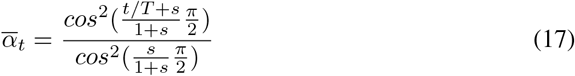

In implementation, *s* was set to 0.008 as recommended in its paper. We also adopted a simplified version of linear noise schedule with Gaussian noise that increased linearly at each time step and added directly to the input data, denoted as *simplified linear noise schedule*.

#### Optimization objective

We employed both noise prediction and reconstruction prediction as the training target to analyse the influence of optimization objective. We also included the combination of standard linear noise schedule in the comparison to better understand the diffusion and denoising process in the model training.

#### Monte Carlo sampling

Similar to EVEscape, we used Monte Carlo sampling to proximate the log likelihood ratio when calculating the mutation fitness prediction. Upon the optimal model architecture, hyper-parameters and noise schedule, we chose the sampling amount of 20,000, 40,000, 60,000 and 80,000 in this experiment to find the best sampling amount with acceptable computation duration.

#### Evaluation Approach

We conducted repeated random sampling to provide a comprehensive evaluation of the model prediction performance in the quantitative comparison experiments. Particularly, we adopted Monte Carlo repeated subsampling and randomly selected 500 mutations each time to measure the fitting degree between experimental data and model prediction, repeating the random sampling 20 times for each model. We followed the same normalization procedures used in previous work [13] to further process both model predictions and experimental data. Given original data *x*_*i*_, along with corresponding mean *µ*_*x*_ and standard deviation *σ*_*x*_ values, they were standardized first, followed by a logistic mapping, and finally a logarithmic transformation to yield the ultimate data processing results *y*_*i*_ as below:

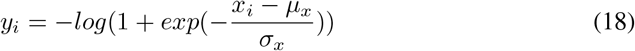

We implemented Mean Squared Error (MSE) as the evaluation metric in each random sampling experiment to directly show the fitting performance of model prediction to the functional experiment data. The standard deviation of 20 random sampling evaluation for each model was further employed as the error bar for box plots and bar charts.

## Data and Code Availability

The data used in this work, including training and evaluation data are cited and credited in the manuscript. Model code and prediction results are uploaded to the paper repository at https://github.com/wuaipinglab/VES.

## Acknowledgements

This work was supported by grants from the Prevention and Control of Emerging and Major Infectious Diseases-National Science and Technology Major Project (No. 2025ZD01901800), the National Natural Science Foundation of China (No. 82550101 & No. 32370703), the Non-profit Central Research Institute Fund of Chinese Academy of Medical Sciences (No. 2023-PT330-01), the CAMS Innovation Fund for Medical Sciences (CIFMS)(No. 2021-I2M-1-061), the Major Project of Guangzhou National Laboratory (No. GZNL2024A01015), the Suzhou Basic and Applied Research Program (Medical and Health) Science and Technology Innovation Project (No. SYW2024065) and the NCTIB Fund for R&D Platform for Cell and Gene Therapy.

## Author Contributions

J.Wu proposed and implemented the model, carried out experiments, evaluated results, and wrote the initial draft of the manuscript. X.Ding proposed and implemented the model, collected and processed data, evaluated results. A.Wu led and supervised the research, provided technical support and advice. All authors reviewed, revised and approved the final version of the manuscript for submission.

---

**Extended Data Figure 1.**
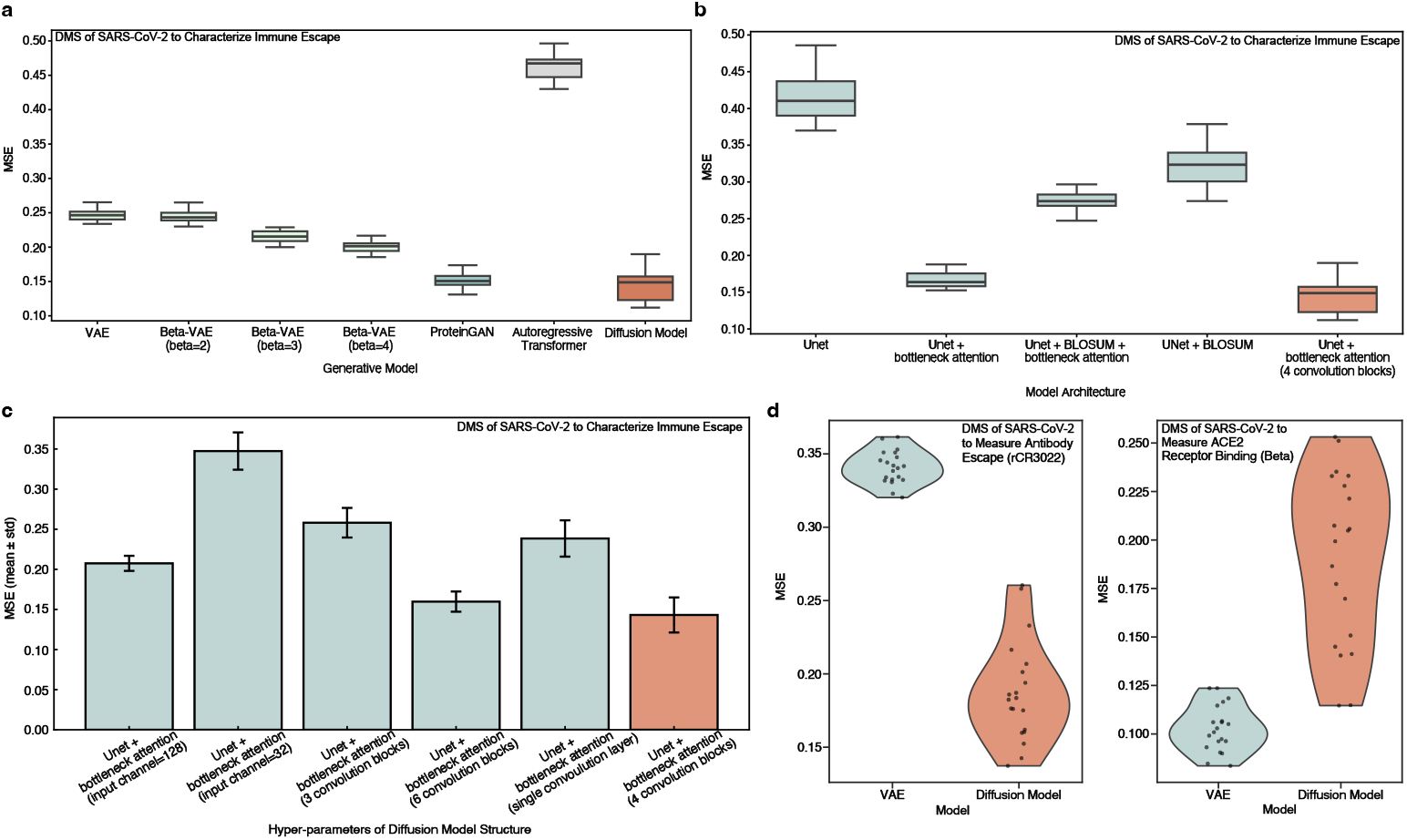
Quantitative benchmarking and ablation analyses of VES on SARS-CoV-2 viruses. **a**, Benchmarking of mutational fitness predictions against experimentally measured immune escape profiles. **b-c**, Ablation study of VES components on DMS of SARS-CoV-2 dataset to characterize immune escape data, including denoising module architecture (b) and hyper-parameters perspectives (c). **d**, Generalization to functional measurements, including DMS of SARS-CoV-2 datasets on antibody escape (rCR3022) and ACE2 receptor binding (Beta), comparing VES with the VAE baseline.

**Extended Data Figure 2.**
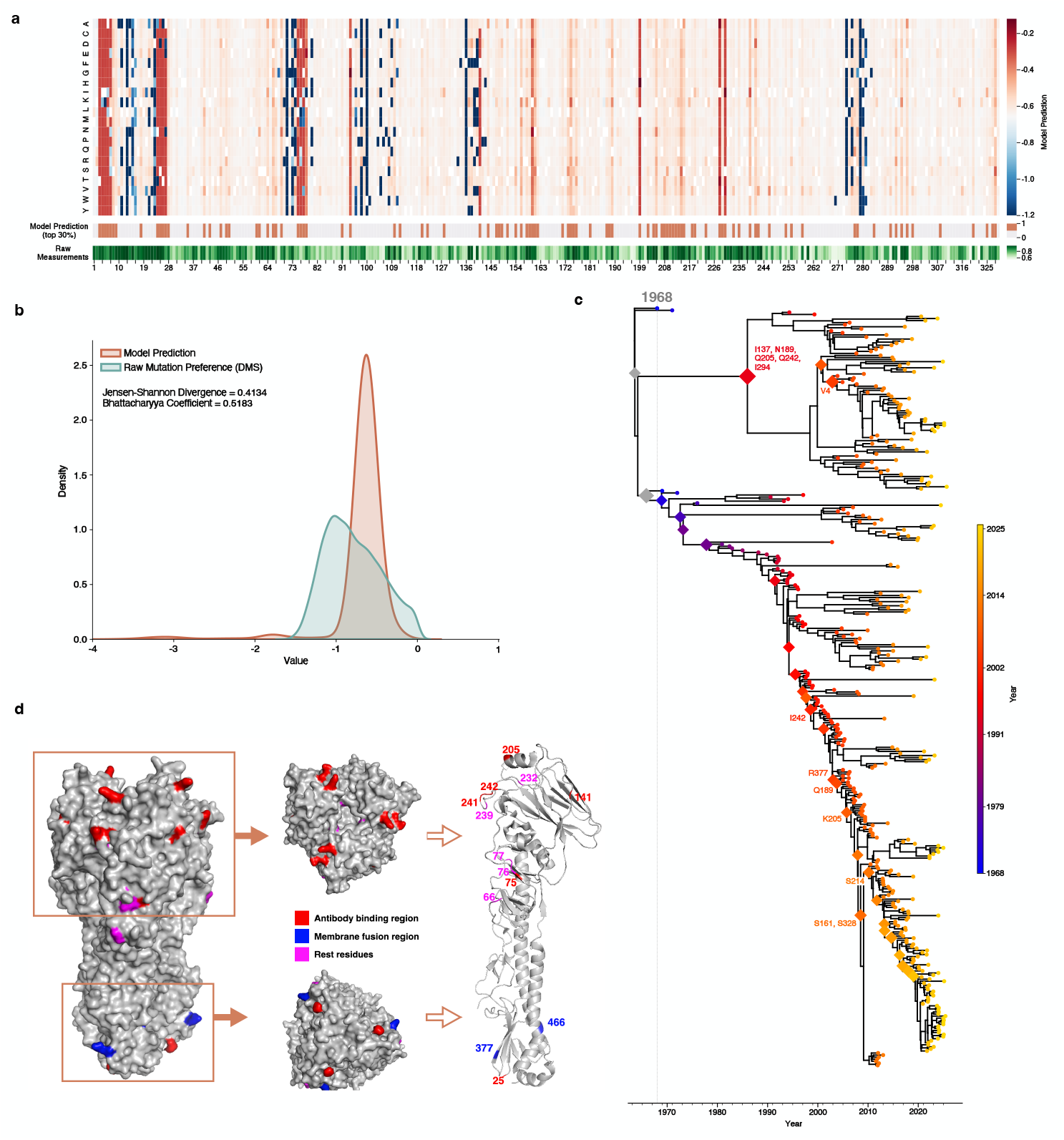
VES maps mutational fitness landscapes and evolutionary trajectories of 8 influenza A/H3 viruses. **a**, Heatmap of site-specific mutational fitness predicted by VES across the HA1 domain. Top-ranked sites prioritized by the model and experimentally validated functional sites are indiated below. **b**, Density distributions of experimentally measured DMS mutation preference values and VES-predicted mutational fitness (Jensen–Shannon divergence of 0.4134, and Bhattacharyya coefficient of 0.5183). **c**, Time-calibrated phylogeny of influenza A/H3 viruses, with high-scoring mutations (top 5% of VES predictions) mapped onto the tree, spanning the major evolutionary stages of seasonal A/H3. **d**, Structural mapping of the top 20 high-fitness sites predicted by VES onto the haemagglutinin (HA) monomer (PDB ID: 4O5N).

**Extended Data Figure 3.**
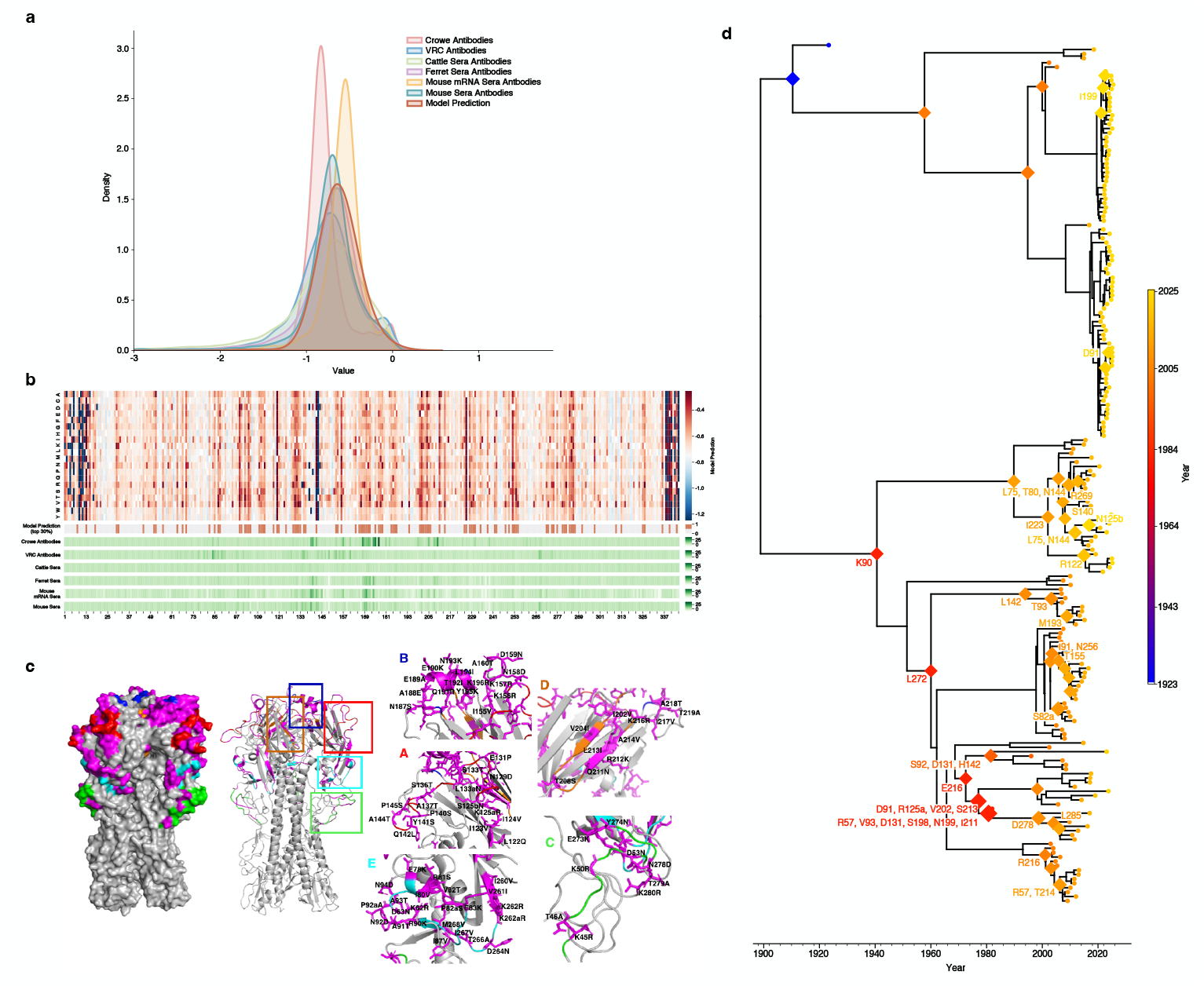
VES predicts antibody escape for influenza A/H5 viruses. **a**, Density distributions of experimentally measured antibody escape profiles (DMS) compared with VES fitness predictions across multiple neutralizing antibodies. **b**, Heatmap of site-specific mutational scores across the HA1 domain under epitope constraints. Residues ranked in the top 30% of model predictions and experimentally validated escape sites are indicated below. **c**, Structural mapping of VES predictions onto the haemagglutinin (HA) monomer, with canonical epitope regions of influenza A/H5 shown in distinct colours and mutational sites in the top 10% of VES predictions highlighted in magenta. **d**, Time-calibrated phylogeny of influenza A/H5 viruses with high-scoring mutations in the epitope regions (top 5% of model predictions) mapped onto the tree.

**Extended Data Figure 4.**
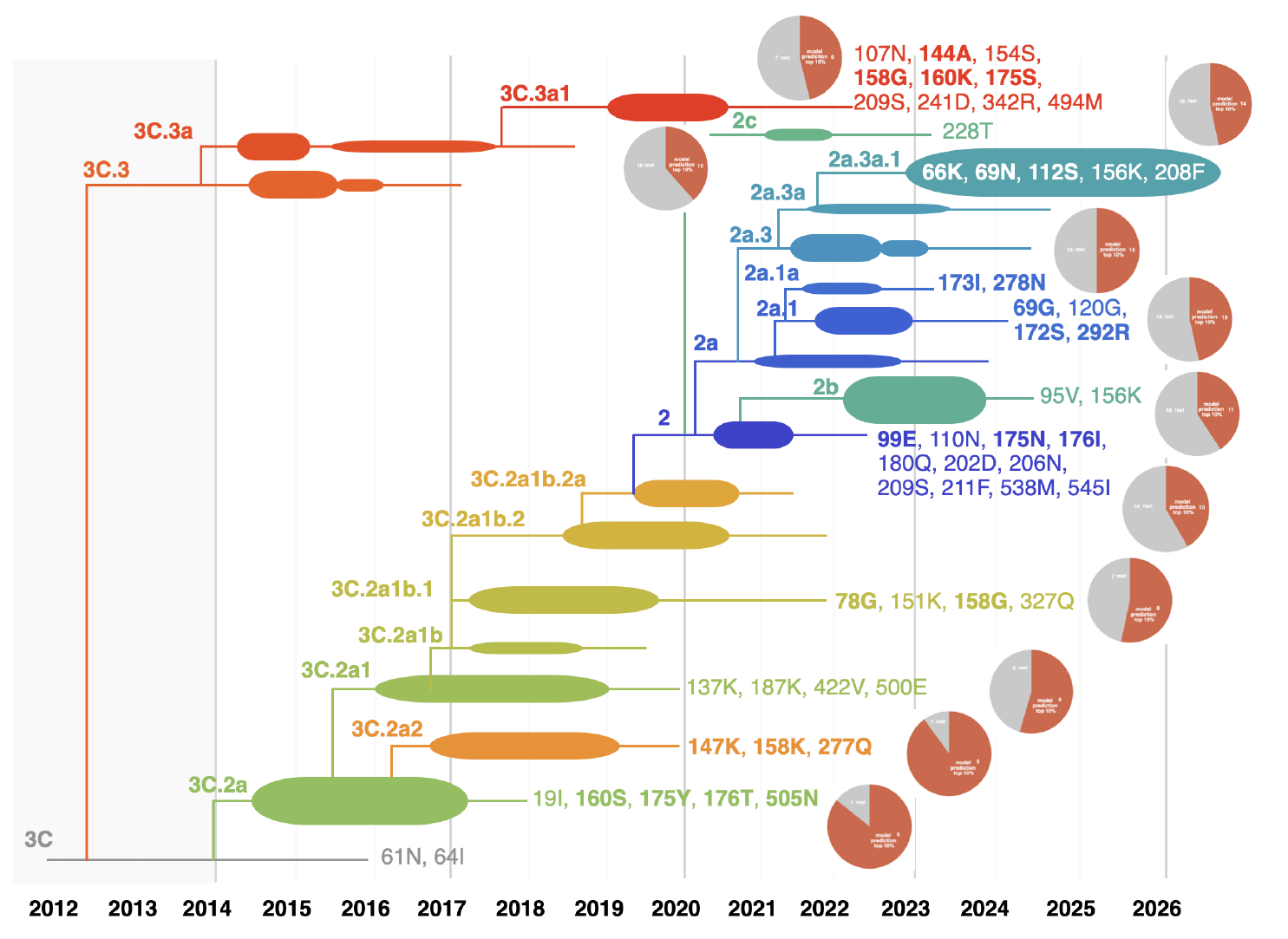
Retrospective analysis on the evolutionary dynamics of influenza A/H3 virus clades since 2014. Sequence data collected before 2014 are adopted to train VES model, and the mutation prediction score is calculated based on it to forecast the evolution of clades that became prevalent after 2014. Clade-defining mutations with top 10% ranked model prediction are shown in bold. The pie charts corresponding to specific clades illustrate the proportion of mutations with top 10% ranked prediction scores (red part) relative to the rest clade-specific mutations (grey part).

**Extended Data Figure 5.**
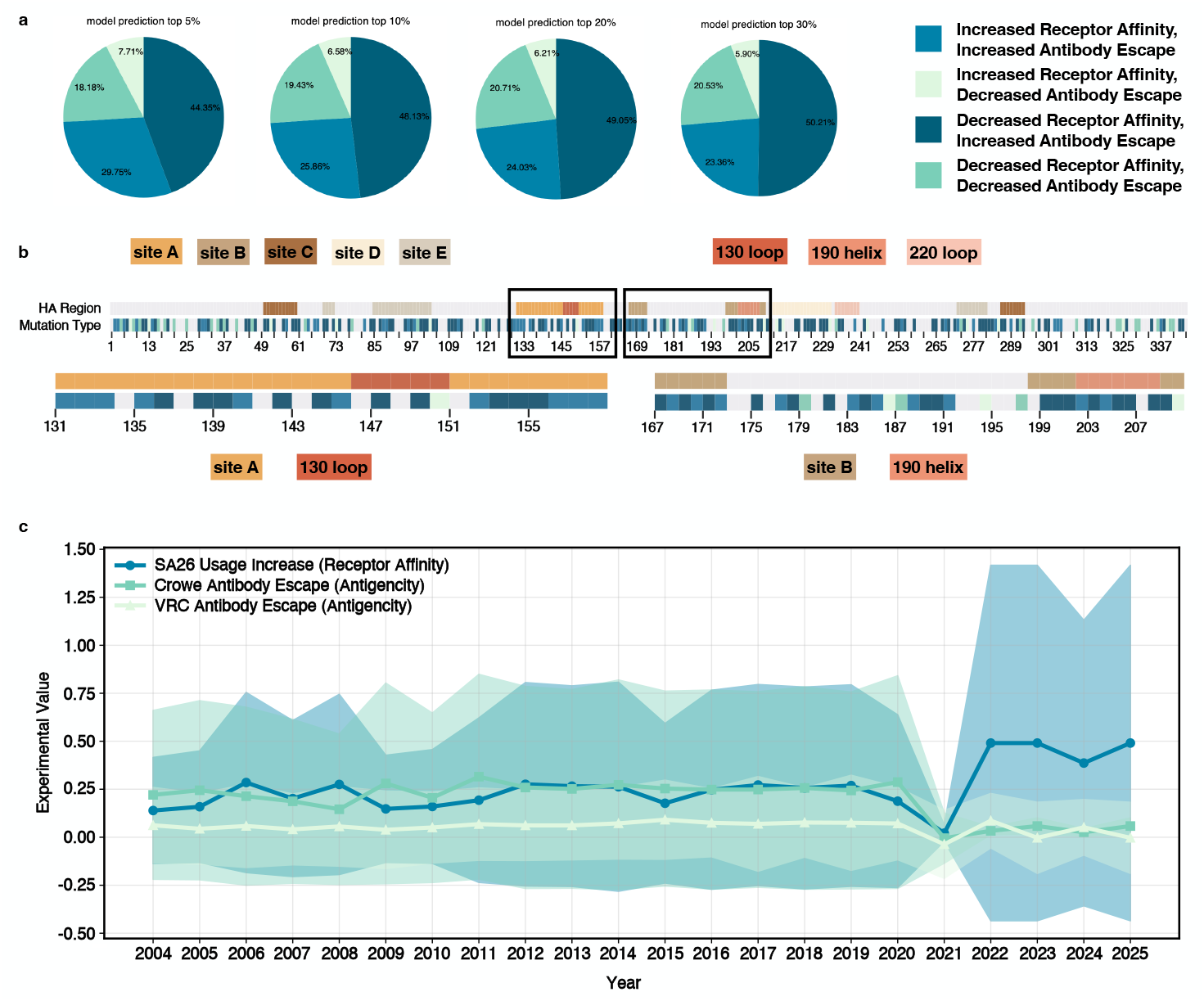
VES discovers the co-evolution pattern of immune escape and receptor binding functions within avian influenza viruses. **a**, Distribution of four co-evolution types for different proportions of model predictions. **b**, Illustration of co-evolution type for sites on HA1 domain. Since there is overlap for antigenic epitope region A and 130 loop sites, as well as antigenic epitope B and 190 helix regions, we magnify the sites in these two overlapping regions to more clearly demonstrate the co-evolution characteristics of functions corresponding to mutations. **c**, Timeline of the co-evolution of antigenicity and receptor affinity from 2004 to 2025. Antigenicity and receptor affinity alternately increased and decreased over time, but both functions experienced significant suppressing in 2021. Subsequently, in 2022, the affinity of avian influenza for human receptors increased significantly, and this high binding affinity continued until 2025, which may partially explain the spread of influenza in recent years.

**Extended Data Table 1:**
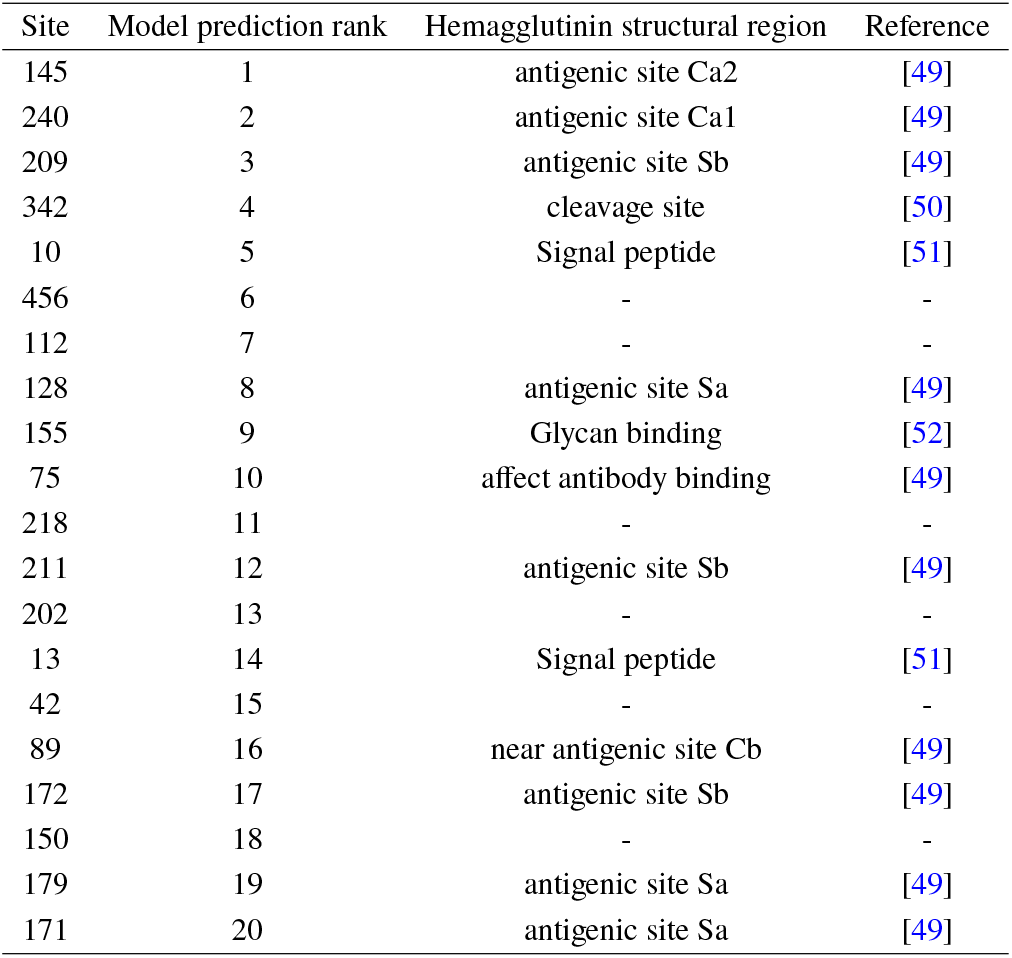
Experimental validation for sites with top 20 ranked model predictions (influenza H1 subtype viruses).

**Extended Data Table 2:** Experimental validation for sites with top 20 ranked model predictions (influenza H3 subtype viruses).

| Site | Model prediction rank | Hemagglutinin structural region | Reference |
| --- | --- | --- | --- |
| 26 | 1 | - | - |
| 3 | 2 | N terminus, antibody binding | [53] |
| 4 | 3 | N terminus, antibody binding | [53] |
| 232 | 4 | - | - |
| 141 | 5 | antigenic site A | [29] |
| 24 | 6 | - | - |
| 6 | 7 | N terminus, antibody binding | [53] |
| 75 | 8 | cluster-difference substitution (H75Q) | [54] |
| 76 | 9 | - | - |
| 239 | 10 | - | - |
| 25 | 11 | cluster-difference substitution (L25I) | [54] |
| 377 | 12 | near the region that influences the pH value required for membrane fusion | [30] |
| 66 | 13 | - | - |
| 5 | 14 | N terminus, antibody binding | [53] |
| 242 | 15 | antigenic site D | [29] |
| 205 | 16 | antigenic site D | [29] |
| 7 | 17 | N terminus, antibody binding | [53] |
| 466 | 18 | near the region that influences the pH value required for membrane fusion | [30] |
| 77 | 19 | - | - |
| 241 | 20 | near antigenic site D | [29] |

